# First experimental evidence for active farming in ambrosia beetles and strong heredity of garden microbiomes

**DOI:** 10.1101/2022.07.27.501732

**Authors:** Janina M.C. Diehl, Vienna Kowallik, Alexander Keller, Peter H. W. Biedermann

## Abstract

Fungal cultivation is a defining feature for advanced agriculture in attine ants and fungus-farming termites. In a third supposedly fungus-farming group, wood-colonizing ambrosia beetles (Curculionidae: Scolytinae and Platypodinae), an experimental proof for the effectiveness of beetle activity for selective promotion of their food fungi over others is lacking and farming has only been assumed based on observations of social and hygienic behaviors.

Here, we experimentally removed mothers and their offspring from young nests of the fruit-tree pinhole borer, *Xyleborinus saxesenii* (Scolytinae). By amplicon sequencing of bacterial and fungal communities of nests with and without beetles we could show that beetles are indeed able to actively shift symbiont communities. Although being consumed, the *Raffaelea* food fungi were more abundant when beetles were present while a weed fungus (*Chaetomium sp*.) as well as overall bacterial diversity were reduced in comparison to nests without beetles. Core symbiont communities were generally low diverse and there were strong signs for vertical transmission not only for the cultivars, but also for secondary symbionts. Our findings verify the existence of active farming, even though the exact mechanisms underlying the selective promotion and/or suppression of symbionts need further investigation.

## Introduction

The cultivation of crops for food is a rare ecological feature, which has evolved only a few times in animals. Apart from humans, the most prominent farmers are fungus-cultivating insect groups specifically one lineage of ants (220 species) and one lineage of termites (330 species) (1,2). Although farming insects are clearly biologically distinct from humans, their farming techniques are remarkably similar, suggesting convergent evolution in response to similar ecological challenges (3). An important example of these shared challenges is the ubiquitous threat of weeds and pathogens for the long-term cultivation of crops. Here, insect farmers evolved a wide variety of strategies to selectively facilitate the growth of their cultivars (1,2,4). These include the sequestration and compartmentalization of gardens (2), usage of antibiotic producing bacteria (5–8) and the active monitoring and behavioural management of fungus garden communities (9–12).

While ants and termites are famous for their pronounced farming practices, this phenomenon likely also evolved in other insect groups. Ambrosia beetles (Coleoptera: Curculionidae: Scolytidae) are a polyphyletic group composed of at least twelve lineages (>3 400 species) of wood-dwelling weevils in the subfamilies Scolytinae and Platypodinae, which are closely associated with symbiotic fungi. Ambrosia beetles carry and introduce these fungal symbionts, which will then grow on the wall of tunnels bored into the xylem of trees and serve as the beetle’s sole food source (13). While the beetles depend on the fungi as food, the fungi are not found outside the beetle-environment demonstrating a strong co-evolutionary history between the partners. Due to this strong interdependent relationship, active management of fungal cultivars has been assumed for ambrosia beetles since the late 19^th^ century (14–16). Ambrosia beetles are ancestrally subsocial (i.e., parental care), with few cases that evolved facultative eusociality and division of labour assumingly in feedback with progressing fungus-farming practices (17). However, because of their enigmatic life within wood, there is very little knowledge on the actual ability of ambrosia beetles to actively promote the growth of their food fungi over others. By contrast, such active farming, which is also a criterion for *advanced agriculture* (sensu 2), has been repeatedly shown for fungus-farming ants and to a lesser degree also termites (9–12,18–21).

The fungal mutualists of Ambrosia beetles, the so-called “ambrosia fungi”, are species-specific (22,23) and typically belong to the ascomycete orders Hypocreales, Ophiostomatales or Microascales (1,24). These yeast-like cultivars produce asexual fruiting structures only in the presence of the beetles and serve as their exclusive food source (25–27). Specifically, they provide their hosts with nutrients (i.e., vitamins, amino acids and sterols (28,29) and essential elements (N, P), which are translocated from the surrounding wood and are strongly concentrated within the fungal tissues the beetles feed on (30). Ambrosia fungus spores are vertically transmitted in the mycetangia (pouch-like structures) or guts of typically female beetles, when they disperse to found new nests and establish their own fungus gardens (31–34). Besides the primary ambrosia fungi, the symbiont community of the beetles consists of other members, such as yeasts (e.g. *Candida, Pichia*), filamentous fungi (e.g. *Penicillium, Aspergillus*) and bacteria (e.g. *Pseudomonas, Orchrobactrum, Bacillus, Enterococcus and Stenotrophomonas*) (22,35–38). There is little knowledge on the role of these “secondary symbionts” for beetle fitness, but the majority of other than ambrosia fungi are assumed to be pathogens or competitors of the beetle-fungus mutualism (24,29,36,38–41).

In some ambrosia beetles, both adults and larvae show cropping behaviour of fungus gardens, which has been hypothesized to have both a weeding and a growth-promoting function, even though it cannot be differentiated from feeding on the fungus (39,41). The reasoning arose from the frequent observation that primary cultivars are quickly overgrown by secondary filamentous fungi following the dispersal of adult beetles from their nests (e.g., (14,22,26,42)) and that nutritional fruiting structures of some ambrosia fungi are only induced in the presence of the beetles (25). Furthermore, there is the assumption of active farming practices in particular species of ambrosia beetles: (i) In *Xyleborus affinis* (Scolytinae: Xyleborini) correlative data showed that cropping behaviour is more commonly expressed when certain fungal symbionts are present (39). (ii) In *Xyleborinus saxesenii* (Scolytinae: Xyleborini) experimental injections of fungal pathogens showed that larvae and adults can actively suppress them by (allo-)grooming and cannibalism of infected individuals (41,43). (iii) Finally, in both *X. affinis* and *X. saxesenii*, an actinomycete, antibiotic-producing bacterial symbiont (*Streptomycetes* sp.) selectively inhibits secondary fungal symbionts but not the ambrosia fungi *in vitro* (6). Nevertheless, all of these indications are no proof for the beetle’s abilities to actively manage fungus-garden microbial communities. A comparison of fungus-garden communities in the presence and absence of their beetle hosts is needed. Given active farming, fungal gardens lacking beetle activity should quickly become dominated by secondary symbionts and may lack the typical succession of the ambrosia fungi (44). Alternatively, cultivars may suppress competitors and pathogens by themselves even in the absence of their host, making active management of beetles unnecessary.

Among ambrosia beetles, the fruit-tree pinhole borer *X. saxesenii* is the behaviourally best studied system to date, because it can be reared and observed in a semi-natural rearing medium (45). The life-cycle starts with the dispersal of an adult female accompanied by two nutritionally important ambrosia fungi (*Raffaelea sulphurea* and *R. canadensis* (Ascomycetes: Ophiostomatales)) within their elytral mycetangia and the gut. These fungi are vertically transmitted to a newly excavated tunnel system in the xylem of a freshly dead tree (34,38,40).

Given that *R. sulphurea* dominates young nests and *R. canadensis* overtakes the garden community in older nests, it has been assumed that the first primarily serves the larvae and the later nurtures the adults (38,44). Nests fail if at least one of these fungal mutualists does not establish or other secondary symbionts (i.e., *Penicillium, Nectria, Chaetomium* and *Aspergillus* species) overgrow gardens initially (6,38,40,46). These antagonists are possibly kept in check by hygienic behaviours of the mother, their larvae and later also adult offspring (6,41,43).

*X. saxesenii* is an inbreeding species, with sib-mating in the natal nest (47). It is among the few known cooperatively breeding beetle species with some adult daughters delaying their dispersal to help their mother in brood care, blocking of the nest entrance, nest maintenance and assumedly fungus farming (13,38,41,43,48,49). Delayed dispersal and reproductive division of labour among adult daughters (i.e., some daughters lay eggs, some do not) is regarded to be selected by these indirect fitness benefits of philopatry, but also by high costs on independent breeding, caused by the low success of establishing an own nest (less than 20% of daughters that reach a suitable substrate can successfully establish the cultivars; (41,50)). Larvae pass three instars (46), whereas the third one is unique for its participation in social nest-hygiene (i.e., the removal of faeces and the ability to suppress fungal pathogens (41)).

Here, we experimentally test for the first time whether farming is indeed part of the ecology/behaviour of ambrosia beetles as assumed since the first behavioural observations of these beetles (14). In particular, we examine if family groups of mothers and larvae actively manage fungus garden communities and thus enhance the longevity/productivity of the fungus garden resource before the first offspring matures. By using *X. saxesenii* as our model we allow adult females to establish nests with fungus gardens in rearing tubes (45). Before eggs are laid, these females are either (i) removed (= *removal group*) and forced to establish a new nest with a fungus garden (= *2*^*nd*^ *attempt group*) or (ii) are allowed to remain in their original nests after they were taken out only briefly to control for the experimental manipulation (= *control group*). From all three treatments, bacterial and fungal communities are determined when the first offspring matures. For fungal communities, newly designed DNA-metabarcoding primers for the large subunit (LSU) are used. These primers will serve as a template for future characterization of ambrosia beetle fungal symbionts as they amplify Ophiostomatales ambrosia fungi, which are typically not picked up by standard ITS barcoding primers (cf. 51).

## Material & Methods

### Beetle rearing and experimental treatments

*X. saxesenii* females were collected in the Steinbachtal near Wuerzburg, Germany (49.767500, 9.896770/49°46’03.0”N 9°53’48.4”E) by using ethanol baited traps (70 % EtOH) in May 2018. After bringing them to the lab they were reared in a sawdust composed “standard medium” in transparent plastic tubes following Biedermann et al. (45). More precisely, these wild-caught adult females (= F0) (that are already mated with brothers before dispersal) were introduced individually into the tubes after rinsing them briefly with 70% EtOH followed by tap water and letting them dry on tissue paper. These founder females immediately start tunnelling when put inside the tubes and 4-7 days later, symbiont growth starts covering the tunnel walls (i.e., “fungus garden”). Fourty dispersing adult female offspring of 11 family groups were used for starting the treatments described in the following.

After all 40 females of the F1 generation successfully established their fungus gardens (7-10 days after nest foundation) three treatment groups were created (see Fig. 1), whereby nests from the same original families were split to *removal* and *control* treatment to control for between-family differences in symbiont communities: (i) *Removal group:* Nests with fungus gardens but no beetles present (N = 20 nests). The solid media was shaken out of the tube and females were removed from the tunnel with a flame-sterilized dissecting needle; if females were not seen from the outside they were removed together with the upmost centimetre of the medium, where they normally reside until the first larvae hatch (41). (ii) *2*^*nd*^ *attempt group:* Subsequently, these females were transferred to the remaining sterile rearing tubes for a second nest foundation (N = 11 nests). Repeated nest founding also occurs in nature (52), but still only 11 out of the 20 second-foundation attempts succeeded and could be used for our analyses. (iii) *Control group:* Nests were treated as in the *removal group* to control for experimental disturbance, but females were put back into their original nests (N = 20 nests). The first two treatments can be compared in a pairwise manner since founding females are the same in both. The *control group* allows to test for possible differences in symbiont communities vectored with 1^st^ and 2^nd^ founding attempts. Additionally, informative metadata, such as family lineage, nest origin, and the exact dates of (a) first introduction to medium, (b) removal of foundress and (c) collection of nest material for amplicon sequencing were recorded.

**Fig. 1.**
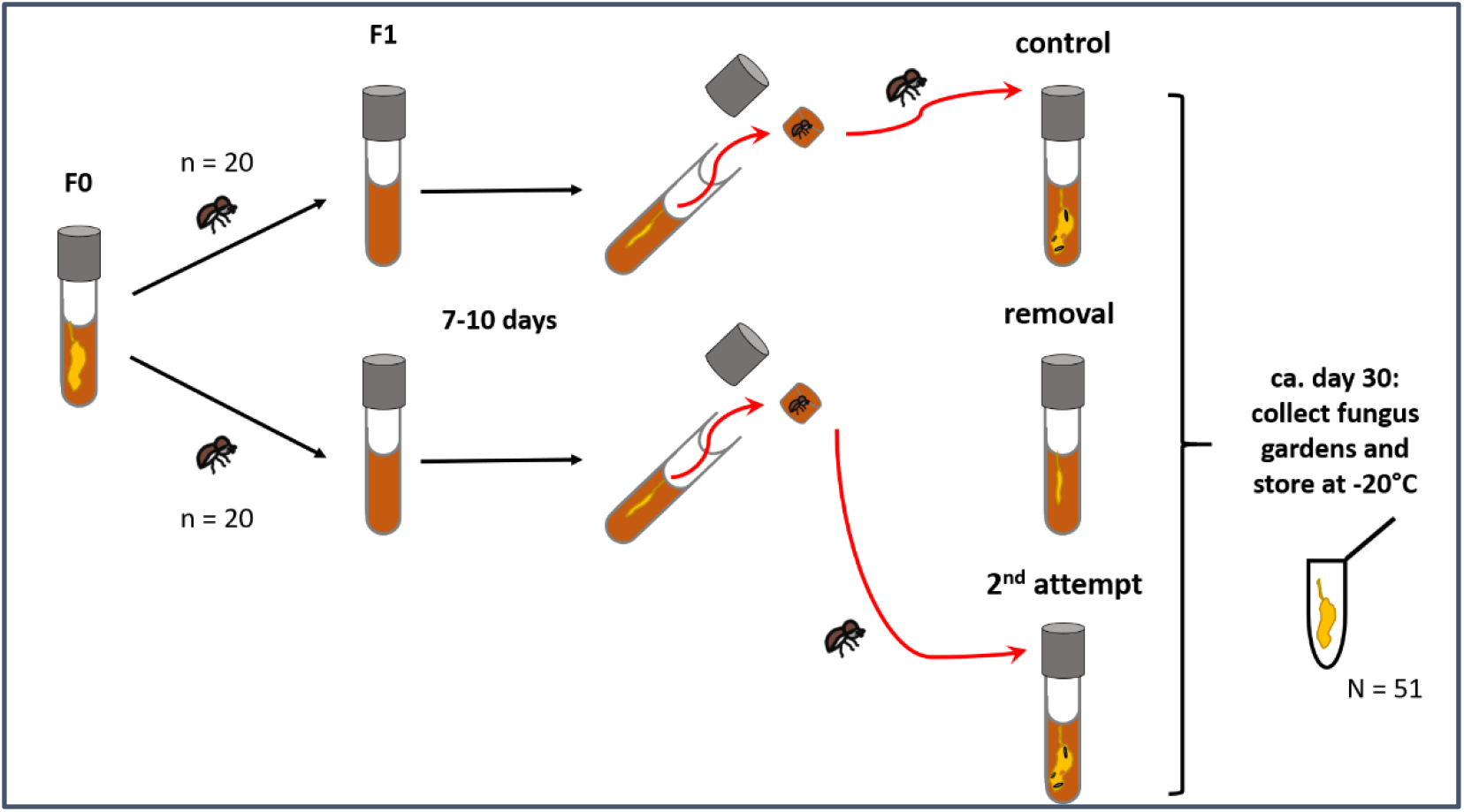
Experimental set-up for the three treatment groups reared in semi-natural media within tubes. Adult sisters from eleven different nests (F0) were equally spread among *control* and *removal group* (F1; each *n* = 20). Seven to ten days later one centimeter of the medium including the foundress was removed and she was either returned immediately to the original nest (*control group*) or was permanently removed from the tube (*removal group*) and introduced immediately into a new rearing tube (*2*^*nd*^ *attempt group*). Eleven of these twenty re-introduced females bred successfully. At around day 30, when first beetles matured, fungus garden samples of all F1 nests were collected and stored at -20°C for bacterial and fungal community sequencing.

### Fungus garden sampling and DNA extraction

When the first females eclosed, nests of all three groups were opened and fungus garden samples were collected. For this purpose, nests were knocked out of the tube under a sterile hood and garden material growing along the gallery walls was sampled with a flame-sterilized spatula (mean ± SD = 96.47 mg ± 34.34). Sampling time did vary between day 19 to day 40 due to variation of development time between nests (mean ± SD = 30.88 days ± 3.09). All samples were mechanically grinded to break up tissues and cell walls using a ceramic bead and mixer mill (Retsch MM400), followed by another step with glass beads (0.1 mm and 0.5 mm) vortexed on a Vortex Genie 2 (see Supplemantary Material in (38)). Afterwards, DNA of all samples was extracted using the ZymoBIOMICS DNA Miniprep Kit (Zymo Research, Germany) according to the manufacturer’s instructions. The isolated DNA samples were stored at -20°C until final amplification and sequencing.

### Library Preparation and Amplicon Sequencing

PCRs were performed in triplicate reactions (each 10 µl) in order to avoid PCR bias. Bacterial 16S rRNA gene libraries were constructed using the dual-indexing strategy described in Kozich et al. (53) using the 515f and 806r primers to amplify amplicon sequences of a mean merged length of 238.18 bp, encompassing the full V4 region (modified from (54)). Conditions for the PCR were as follows: initial denaturation at 95°C for 4 min, 30 cycles of denaturation at 95°C for 40 s, annealing at 55°C for 30 s and elongation at 72°C for 1 min; followed by a final extension step at 72°C for 5 min. Sample-specific labelling was achieved by assigning each sample to a different forward/reverse index combination.

For sequencing of the fungal communities, the LSU (28S) rRNA gene libraries (mean merged length of 276.74 bp) were constructed similarly from the same samples to amplify the large subunit (LSU) region. Again, adapters and dual-indices were incorporated directly into the PCR primers. Conditions for this PCR with the new, self-designed and dual-index primers of LIC15R and nu-LSU-355-3’ (described in Supplementary Material in (38)) were as follows: initial denaturation at 98°C for 30 sec, 35 cycles of denaturation at 98°C for 30 s, annealing at 55°C for 30 s and elongation at 72°C for 15 sec; followed by a final extension step at 72°C for 10 min.

After both PCRs, triplicate reactions of each sample were combined per marker and further processed as described in Kozich et al. (53), including between-sample normalization using the SequalPrep™ Normalization Plate Kit (Invitrogen GmbH, Darmstadt, Germany) and pooling of 96 samples. The pools were cleaned-up with the AMPure Beads Purification (Agilent Technologies, Inc. Santa Clara, CA, USA), quality controlled using a Bioanalyzer High Sensitivity DNA Chip (Agilent Technologies, Santa Clara, CA, USA) and quantified with the dsDNA High Sensitivity Assay (Life Technologies GmbH, Darmstadt, Germany). Afterwards, pools were combined to a single library pool containing 384 samples in total. This library was diluted to 8 pM, denatured and spiked with 5% Phix Control Kit v3 (Illumina Inc., San Diego, CA, USA) according to the Sample Preparation Guide (llumina Inc. 2013). Sequencing was performed on a Illumina MiSeq using 2 × 250 cycles v2 chemistry with each marker on a separate chip (Illumina Inc., San Diego, CA, USA). See Supplementary Material for further methodology of sequencing controls and details on bioinformatics processing.

### Statistical analysis of molecular data

All statistical analyses and visualisation of the sequence output were performed in RStudio (version 1.4.1106) with R version 4.0.5 (55) using the phyloseq package ((56); see GitHub repository for information on the bioinformatic processing and R-script).

After filtering Chloroplast genes and amplicon sequence variants, that were only identified to domain level, we excluded control samples, as well as samples with a read number < 500. For the final analysis 28 samples with an average of 15,011.72 reads for 16S sequences [min. 793 reads; max. 44924 reads] and 69 ASVs (amplicon sequence variants, (57)) were included. Microbial composition of the bacteria was studied up to the genus level and their relative abundance. For the LSU, 51 samples with an average of 17,344.02 reads [min. 1087 reads; max. 39932 reads] and 202 ASVs were included in the analyses. Fungal composition was studied up to the species level and their relative abundance.

For the analysis of the alpha diversity we rarefied the sequence reads of all samples to a total of 2000 reads per sample and tested the observed estimate of taxa richness and Shannon diversity index (‘microbiome’ package: (58)). Rarefying removed six further samples and 15 ASVs from the 16S dataset and two samples, as well as, 23 ASVs from the LSU dataset.

#### Testing for active farming between sisters: *removal group* vs. *control group*

To test for the influence of the beetle’s presence on the fungus garden microbiome, we compared the microbial community between nests in the *control group* and the *removal group*. Alpha diversity of the rarefied samples was explored by plotting Observed species richness (OR) and Shannon’s diversity index (SDI). For the bacterial alpha diversity, we ran a generalized linear model (GLM) with gamma family and log link function on the SDI to test the influence of the treatment (*‘control’* vs *‘removal’*) and ‘family lineage’. A GLM with normal distribution best fitted the fungal diversity data. For OR (number of observed ASV’s) we applied the same model. The package ‘ggplot2’ (59) was used to build the figures of alpha diversity (SDI/OR).

To visualize differences in composition (beta diversity), non-metric multidimensional scaling (NMDS, ‘phyloseq’ package: (56)) was used on Bray-Curtis dissimilarity matrices derived from proportion transformed data, which consider presence/absence as well as abundance of ASVs (60). To compare the microbial communities between the treatments and ‘family lineage’, we performed a permutational ANOVA test (PERMANOVA) on Bray-Curtis distance matrices of the proportion data using the R package ‘vegan’ (61). The homogeneity of multivariate dispersions was tested with betadisper() and distance structures of the bacterial and fungal data (‘vegan’ package: (61)) were applied on each the ‘treatment’ and ‘family lineage variables’. Taxa composition barplots [agglomerated to ‘genus’ (bacteria) or ‘species’ level (fungi)] faceted by lineage and heatmaps of fungal and bacterial communities were built for visualization. As ASVs can represent biological variance between microbial strains of the same species, we wanted to test for specific associations or variations between family lineages and therefore plotted an additional bar graph of the highly abundant strains of core species (ASVs >0.5%) from the *control* and *removal group*.

#### Testing for active farming by controlling foundress identity: *removal* group vs. *2*^*nd*^ *attempt* group

Comparing the *2*^*nd*^ *attempt* with the *removal group* gave us the opportunity to directly study the effect of the beetle’s presence, by controlling for between individual differences in microbial symbionts of founding females (because the same females were used in both groups). Eleven of twenty females failed in their 2^nd^ founding attempt. In addition to the GLMs, NMDSs and PERMANOVAs and figures we used in the previous analysis, we additionally ran a linear model (LM) applying logistic transformation on the SDI of the bacteria to test the influence of the treatment (*‘2*^*nd*^ *attempt’* vs *‘removal’*) by controlling for ‘family lineage’.

#### The responses of core bacterial and fungal taxa

For gaining deeper insights into differences/changes of abundant core taxa, we agglomerated the same taxa (‘genus’ level for bacteria and ‘species’ level for fungi) of the compositional phyloseq object and extracted the individual core taxa. We ran LMs to test whether relative abundances of core taxa differed between treatments and ‘family lineage’. Precisely, we compared the logistic transformed relative abundances of the two ambrosia fungi, *R. sulphurea* and *R. canadensis*, the commensal fungus *C. globosum* and the bacterium *Pseudoxanthomonas* for the between the treatments and familiar lineages. The abundance data of *Wolbachia* was subjected to a Tukey transformation to transform the response variable more towards a normal distribution before being used in a LM with the same variables (‘rcompanion’ package: (62)). Relative abundance boxplots of core taxa were built for both bacteria and fungi.

#### Effects of the microbial community of nests in the *removal group* on the success rate of the same foundresses during the 2^nd^ foundation attempt

The *removal* treatment subset provided us with the possibility to compare nests of foundresses being successful in their *2*^*nd*^ *attempt* (‘*successful*’) with those that failed to found a 2^nd^ brood (‘*failed*’). To examine this, we performed two PERMANOVAs on Bray-Curtis distance matrices of the relative abundances data for each bacterial and fungal communities and tested the homogeneity as previously described. Since core taxa were of particular interest to us, we compared the logistically transformed relative abundances of core taxa between failed and successful 2^nd^ attempts using a LM and plotted them with boxplots.

## Results

### Detected taxa in bacterial and fungal datasets

Altogether ten bacterial phyla were detected across samples. Among these, Gammaproteobacteria, Alphaproteobacteria and Actinobacteria were most abundant and accounted for approx. 90% of total sequences (Fig. 2; Tab. S1). Gammaproteobacteria comprised ASVs of *Pseudoxanthomonas* (mean + SD = 59.93% ± 44.28 RA), *Erwinia* (7.83% ± 22.86) and *Acinetobacter* (0.14% ± 0.85). Alphaproteobacteria were dominated by *Wolbachia* (Alphaproteobacteria) (28.11% ± 43.28) and *Ochrobactrum* (Alphaproteobacteria) (2.67% ± 11.62) and Actinobacteria by *Microbacterium* (0.63% ± 1.16). All abundant bacteria showed to be relatively equally distributed across samples while the endosymbiont *Wolbachia* was only highly dominant in few samples/lineages. Bacteroidetes, Firmicutes, Acidobacteria, Chloroflexi, Verrucomicrobia, Planctomycetes and Fusobacteria were detected in abundances <0.5% of total reads.

**Fig. 2.**
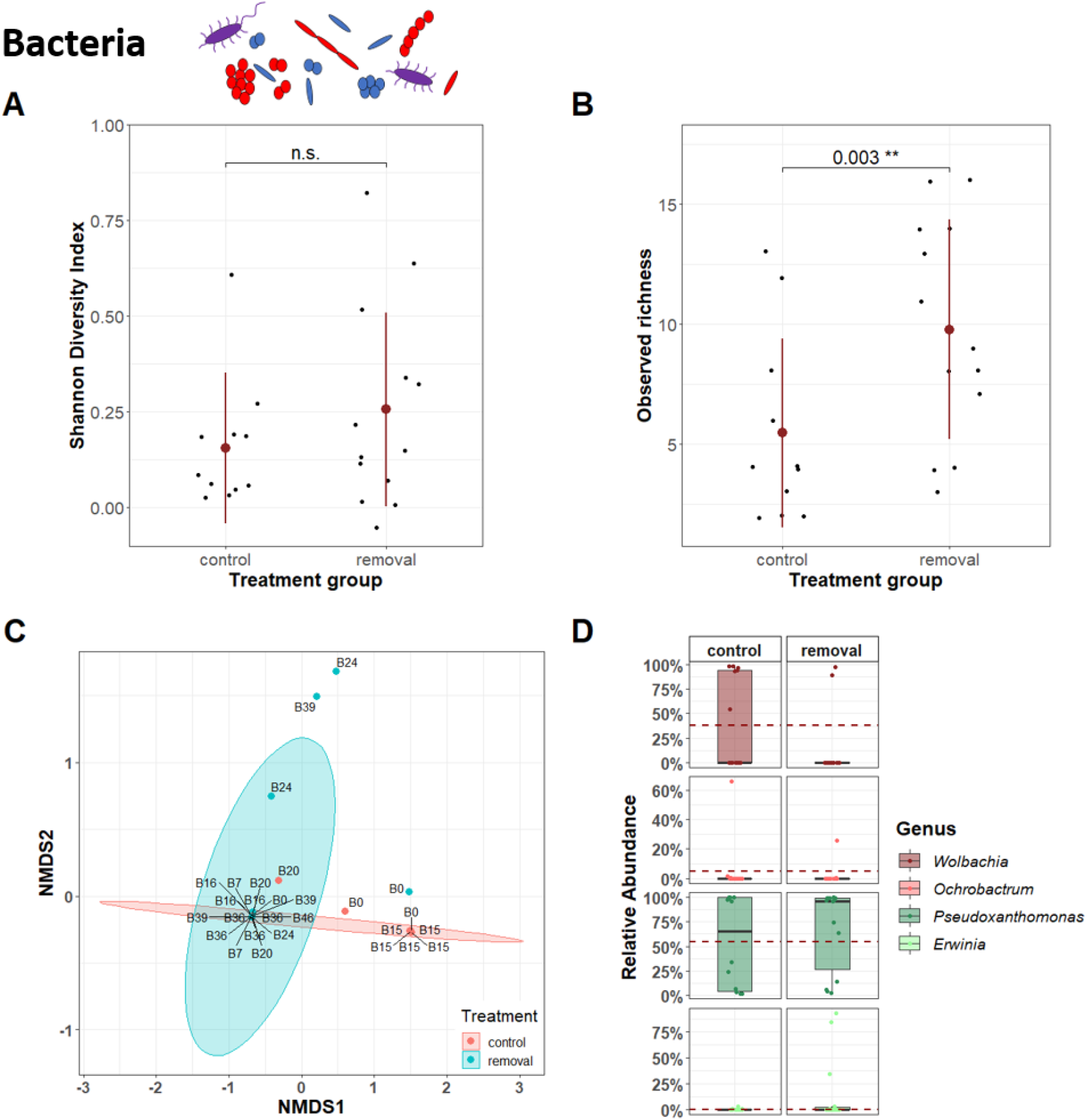
Effect of beetle removal on bacterial diversity, composition and abundance. (A) Bacterial SDI did not differ between in *removal* and *control group* (each *n* = 20 nests; GLM: χ^2^ = 1.57, *p* = 0.210; plot shows mean and standard deviation). (B) Bacterial OR was higher when beetles were removed (χ^2^ = 8.81, *p* = 0.003; plot shows mean and standard deviation). (C) Visualization of compositional differences between bacterial communities of nests in *removal* and *control group* (NMDS on Bray-Curtis dissimilarity: stress = 0.004; ellipses represent 95% confidence intervals; labels represent family lineage of samples). PERMANOVA verified that bacterial communities are significantly affected by beetle presence (*R*^*2*^ = 0.066, *p* = 0.038). (D) Comparisons of relative abundances of core bacterial taxa did not reveal any significant differences between treatments (boxplots represent median with its interquartile range and whiskers; red dashed line shows the mean relative abundance of the *’control group’*).

The analyses of the fungal dataset yielded exclusively ASVs of the phylum Ascomycota. The most abundant species was *C. globosum* (Chaetomiaceae) with a relative abundance of 24.19% ± 34.18 (mean ± SD) followed by the Ophiostomatales ambrosia fungi of *X. saxesenii, Raffaelea sulphurea* (18.46% ± 26.89) and *R. canadensis* (4.17% ± 8.23). Other species with a relative abundance of >0.5% in some samples were *Acremonium biseptum* (Bionectriaceae; 2.48% ± 13.75) and *Penicillium commune* (Trichocomaceae, 0.96% ± 5.23) (Tab. S2). A closer look on our own fungal mock communities revealed that the newly designed primers could distinguish between the two ambrosia fungi *R. canadensis* and *R sulphurea*. Moreover, additional fungi in the orders Eurotiales, Sordariales, Hypocreales, Capnodiales, Onygenales and Dothideales were successfully amplified, but yeasts in the Saccharomycetales order (e.g. *Pichia sp*., *Candida sp*.) could not be differentiated by our approach (cf. (38); Supplementary Figure 5).

### Testing for active farming

#### Bacterial symbiont communities

Comparisons of alpha diversity of bacterial communities between treatment groups showed observed bacterial richness (OR) was significantly higher in the ambrosia gardens without beetles (*control* vs. *removal:* GLM: *χ*^*2*^ = 8.81, *p* = 0.003; Fig. 2B, Tab. 1; *removal* vs. *2*^*nd*^ *attempt*: GLM: *χ*^*2*^ = 4.51, *p* = 0.034; Fig. S3, Tab. 1), but Shannon’s diversity (SDI) did not reveal differences (*control* vs. *removal:* GLM: *χ*^*2*^ = 1.57, *p* = 0.210; Fig. 2A, Tab. 1; *removal* vs. *2*^*nd*^ *attempt*: LM: *F* = 0.406, *p* = 0.538; Fig. S3, Tab. 1). There was also a significant difference in bacterial beta diversity (i.e., the turnover of taxa) between groups (*control* vs. *removal:* PERMANOVA: *R*^*2*^ = 0.066, *p* = 0.038, Betadisper: *F* = 0.771, *p* = 0.388; Tab. 1; *removal* vs. *2*^*nd*^ *attempt*: PERMANOVA: *R*^*2*^ = 0.042, *p* = 0.169, Betadisper: *F* = 0.478, *p* = 0.497; Tab. 1), which was also visible in the NMDS plot (Fig. 2C, Fig. S3). Comparing the relative abundances of core bacterial taxa, neither the two most abundant taxa *Wolbachia* (*control* vs. *removal:* LM: *F* = 2.41, *p* = 0.138; *removal* vs. *2*^*nd*^ *attempt*: LM: *F* = 0.409, *p* = 0.534) and *Pseudoxanthomonas* (*control* vs. *removal:* LM: *F* = 0.177, *p* = 0.679; *removal* vs. *2*^*nd*^ *attempt*: LM: *F* = 0.163, *p* = 0.693) nor any of the other core bacterial ASVs showed a significant response to beetle presence (Fig. 2D, S3, Tab. S3).

**Tab 1.**
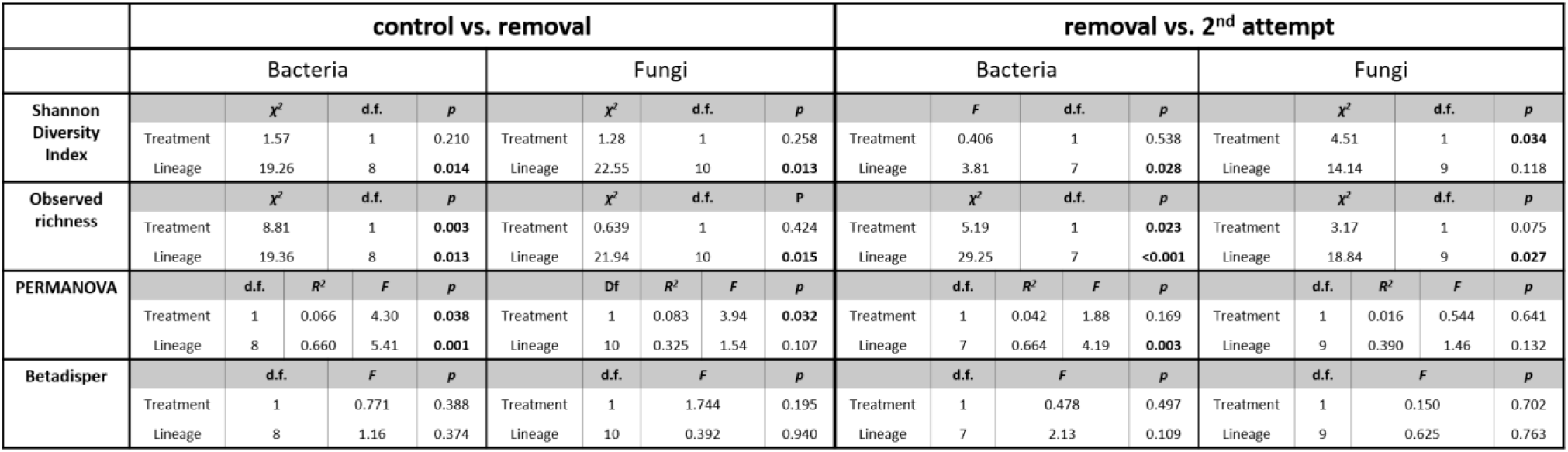
Statistical output of alpha and beta diversity analyses in both bacterial and fungal communities. Both comparisons of *control* vs. *removal* groups and *removal* vs. *2*^*nd*^ *attempt* group are given. Statistically significant results are highlighted (*p* < 0.05).

#### Fungal symbiont communities

Comparisons of alpha diversity of fungal communities between treatment groups did not reveal differences neither for SDI (GLM: *χ*^*2*^ = 1.28, *p* = 0.258; Fig. 3A) nor OR (*χ*^*2*^ = 0.639, *p* = 0.424; Fig. 3B, Tab. 1) when comparing *control* with *removal group*. However, when comparing *removal* with *2*^*nd*^ *attempt group* directly, SDI was significantly higher in the nests without beetles (*χ*^*2*^ = 4.51, *p* = 0.034; Fig. S3; Tab. 1) and also OR showed this tendency (*χ*^*2*^ = 3.17, *p* = 0.075; Fig. S3, Tab. 1). As observed in bacteria, there was also a significant difference in fungal beta diversity between *control* and *removal groups* (PERMANOVA: *R*^*2*^ = 0.083, *p* = 0.032; Betadisper: *F* = 0.771, *p* = 0.388; Tab. 1), but not between *removal* and *2*^*nd*^ *attempt groups* (PERMANOVA: *R*^*2*^ = 0.016, *p* = 0.641; Betadisper: *F* = 0.150, *p* = 0.702; Tab. 1). The NMDS plots displayed strong overlap of the fungal communities (Fig. 3C, S3).

**Fig. 3.**
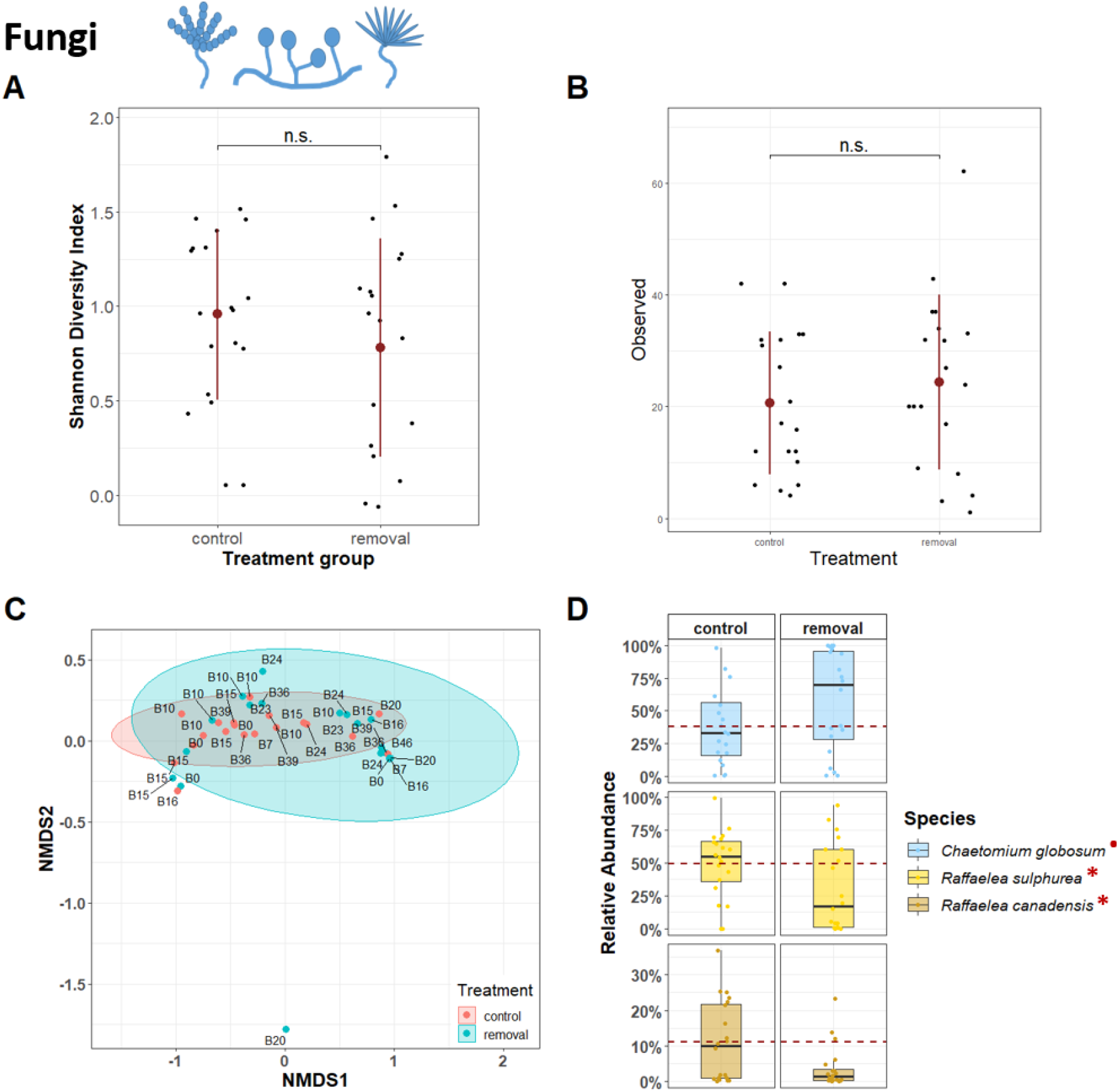
Effect of beetle removal on fungal diversity, composition and abundance. (A, B) Fungal SDI and OR did not differ between in *removal* and *control group* (each *n* = 20 nests; SDI: GLM: χ^2^ = 1.28, *p* = 0.258; OR: χ^2^ = 0.639, *p* = 0.424; plot shows mean and standard deviation). (C) Visualization of compositional differences between fungal communities of nests in *removal* and *control group* (NMDS: stress = 0.034; ellipses represent 95% confidence intervals; labels represent family lineage of samples). PERMANOVA verified that fungal communities are significantly affected by beetle presence (*R*^*2*^ = 0.083, *p* = 0.032). (D) Comparisons of relative abundances of core fungal taxa revealed significant reduction of food fungi, *R. canadensis* (LM: *F* = 5.18, *p* = 0.031) and *R. sulphurea* (*F* = 4.97, *p* = 0.034), in nests without beetles, whereas relative abundance of *C. globosum* tended to increase in these nests (*F* = 3.37, *p* = 0.077; * - *p* < 0.05; • *p* < 0.1. (boxplots represent median with its interquartile range and whiskers; red dashed line shows the mean relative abundance of the ‘*control group’*).

Comparing the relative abundances of core taxa between the treatments revealed strong responses to beetle presence (Fig. 3D; Tab. S3). While *C. globosum* tended to be more abundant in the absence of beetles (LM: *F* = 3.37, *p* = 0.077), both ambrosia fungi, *R. sulphurea* and *R. canadensis*, were significantly more abundant in the presence of beetles (*R. canadensis*: *F* = 5.18, *p* = 0.031; *R. sulphurea*: *F* = 4.97, *p* = 0.034). There was no significant difference between *removal* and *2*^*nd*^ *attempt* groups for *C. globosum* (*F* = 0.092, *p* = 0.765) and *R. sulphurea* (*F* = 1.46, *p* = 0.241), but *R. canadensis* tended to have a higher relative abundance when beetles were present (*F* = 4.30, *p* = 0.051).

#### The effect of the microbial community on the success rate of a 2^nd^ foundation attempt

We found no significant effect of neither the overall bacterial community (PERMANOVA: *R*^*2*^ = 0.071, *p* = 0.573, Betadisper: *F* = 0.021, *p* = 0.889) nor the relative abundance of the core bacterial taxa (*Erwinia*: LM: *F* = 0.641, *p* = 0.439; *Pseudoxanthomonas*: *F* = 0.044, *p* = 0.837; *Wolbachia*: *F* = 2.46, *p* = 0.143) of *removal* nests on the success rate of 2^nd^ founding attempts (Fig. S2). There was also only a trend for an effect of the overall fungal community (PERMANOVA: *R*^*2*^ = 0.091, *p* = 0.164, Betadisper: *F* = 0.2.11, *p* = 0.164) of *removal* nests on the success rate of 2^nd^ founding attempts. However, when testing the effect of the relative abundance of core fungal taxa, *C. globosum* turned out to be significantly less abundant in *removal* nests of successful foundresses (LM: *F* = 5.51, *p* = 0.031). On the other hand, at least one ambrosia fungus tended to be more abundant in these nests (*R. canadensis*: *F* = 2.85, *p* = 0.108; *R. sulphurea*: *F* = 2.10, *p* = 0.164), indicating effects of at least *C. globosum* and possibly also *R. canadensis* on the success rate of foundresses.

#### Heredity of the symbiont community

Overall there were strong signatures for heredity (i.e., vertical transmission) of symbiont communities over generations as seen by the effect of ‘family lineage’ in almost every model (Tab. 1). By including the variable ‘family lineage’ in our models we were able to control for this factor (Fig. S6, Tab. 1). Interestingly we found two strains of *R. canadensis* in our data set, which never appeared together in one sample or within samples of the same family lineage (Fig. S7).

## Discussion

Farming is a complex behaviour that evolved several times across the animal kingdom and is strongly affecting the ecology and evolution of the farmers and their cultivars (3). This study aimed for the first time to test if farming (i.e., the active behavioural management of symbiont communities) is present in ambrosia beetles. Latter is known from other fungus farming insects (1,2) and has been assumed since the first descriptions of ambrosia beetles without proper proof (e.g. (14)). By investigating the effect of beetle presence or removal on the composition of microbial communities, we were able to show that active farming is indeed present in our study species, the fruit-tree pinhole borer *Xyleborinus saxesenii* (Scolytinae, Xyleborini), even if their managing abilities are not as effective as in fungus-farming ants and termites (2).

### To farm or not to farm – the debate of active fungus-farming in ambrosia beetles

Apart from humans, attine ants, fungus-farming termites and ambrosia beetles were defined as agriculturists even though active farming, as one defining feature of agriculture, has never been proven for ambrosia beetles (2). Our findings show that active behavioural farming of cultivars is present in *X. saxesenii*, supporting this premature claim. Firstly, relative abundances of the nutritional ambrosia fungi (two *Raffaelea* spp.) significantly increased in the presence of *X. saxesenii* foundresses and their first immature/larval offspring. Secondly, bacterial alpha diversity and the relative abundance of a fungal competitor of the ambrosia fungi (*Chaetomium* sp.) were both significantly reduced by the presence of foundresses and their immature offspring. Together this confirms old assumptions by Hubbard (14) and many others (e.g. (2,29,63)) that ambrosia beetle species (or at least *X. saxesenii*) can actively manage its fungus-garden communities to some degree.

### The presence of beetles affects symbiont communities

An important finding of our study was that feeding mothers and immatures were not reducing the amount of their *Raffaelea* food fungi, but instead relative fungal abundances increased in their presence (Fig. 3D). Fungus-farming ants and termites are known to directly influence growth of their cultivars, for example by inducing nutritional structures, which has been also shown for one ambrosia beetle species (*Anisandrus dispar* with its *Ambrosiella* symbiont; (2,25)). Furthermore, there is tremendous evidence in all insect farmers for indirect promotion of cultivars by direct suppression of fungal competitors and pathogens. In ambrosia beetles, *X. saxesenii* larvae can suppress the growth of a fungal pathogen (41). Moreover, adult females of the same species upregulated allogrooming and cannibalism (i.e., removal of corpses) after experimental injection of a pathogenic *Aspergillus* sp. in their nests, which effectively reduced its spore loads (43). Such effects may explain the reduced diversity and abundances of non-beneficial secondary symbionts (bacteria and *Chaetomium* sp.) in the presence of mothers and immatures we found here.

Known hygienic behaviors of ambrosia beetles such as the compartmentalization of fungus gardens, allo-grooming and removal of infected nestmates have all been observed before in *X. saxesenii* (and other ambrosia beetles), but their effects on symbiont communities have so far been unknown (1,13,17). Our results clearly show that their effectiveness are by no means comparable to the ones of farming ants and termites that maintain more or less monocultures of their fungal cultivars over several years and including behaviours such as pathogen alarm (2,64). Ambrosia beetles, on the other hand, typically live in small subsocial to cooperatively breeding societies (with maximum 100 individuals in *X. saxesenii*), with relatively short durability (max. 2 years in *X. saxesenii*) (2). Nevertheless, it is possible that the effectiveness of farming was underestimated by our study because we only tested the effect of mothers and their larvae. A stronger effect can be expected when ambrosia beetle galleries are sampled at a later stage, during the presence of mature daughters that delay dispersal and display various hygienic behaviors (41).

### Putative mechanisms for regulating symbiont communities

The exact mechanisms underlying the suppression of secondary symbionts remain unknown, but it is likely that beetle secretions and/or antibiotic-producing bacteria are playing a role (2,5). Yet unknown selective secretions nurture fungal spores during hibernation and dispersal within ambrosia beetle’s cuticular mycetangia (65) and it is possible that these selective secretions are also released in the nest environment. Against this hypothesis speaks the observation that the mycetangial glands are only active before and during adult female dispersal, but are reduced soon after ejection of fungal spores and establishment of a fungus garden (65,66). In *X. saxesenii* release from selective secretions may be happening through the gut, however, because adult females in this species use the gut as a second mycetangium for transmitting *R. sulphurea* (34). Similarly, both termite faeces and saliva have anti-microbial/-fungal activity and have been implicated to reduce the microbial load when applied on dead nestmates (67,68).

Another already proven source of selective compounds are symbiotic bacteria. A *Streptomyces griseus* (Actinobacteria) strain with selective inhibition of secondary fungi (but not the cultivars) has been isolated from both *X. saxesenii* and *Xyleborus affinis* (6). Here we could not detect this strain, but possibly other symbionts provide similar functions, given that screens for antibiotic-producing bacteria in related bark beetles revealed lots of taxa of which some are also present in our community (e.g. *Microbacterium, Pseudoxanthomonas*; (5,69–71). Moreover, both fungus-growing ants and termites are known to use antibiotic-producing bacteria to protect their fungal gardens from invaders (8,72). Workers in both groups can perceive pathogen presence and apply chemical defenses locally (9,21).

### The bacterial community and potential functions

In ambrosia beetles, so far, most descriptive and experimental studies focused on the fungal symbionts (but see (38,73–75)). However, given that ambrosia beetle communities resemble those of fungus-farming ants and termites (73), where bacteria are known to play essential roles (e.g. (8,76–78)), more functional studies on these symbionts are crucially needed.

The overall number of core bacteria was surprisingly small and common taxa belonged to the Actinobacteria, the Alphaproteobacteria and the Gammaproteobacteria, with *Pseudoxanthomonas, Acinetobacter, Erwinia, Ochrobactrum, Microbacterium* and *Wolbachia* being the most abundant phylotypes. This result closely resembles communities found by other studies for *X. saxesenii* and other ambrosia beetles (38,73,75). One of the most abundant genera in our samples was *Pseudoxanthomonas*. This bacterium had been detected in the gut of *X. saxesenii* by Fabig (79) before, but only in a low relative abundance (1.2%). Other studies repeatedly isolated this bacterium from gut samples of bark beetles and profiled it as a cellulolytic bacterium, similar to *Ochrobactrum*, able to produce an array of cellulolytic-xylanolytic enzymes (74,80–82). *Erwinia*, on the other hand, may be able to fix atmospheric nitrogen (83,84), which may profit the farmers in this nitrogen-deficient wood substrate.

*Wolbachia* infections were found in four out of eleven different family lineages that were also passed on between F0 and F1 generations. This and our previous study (38) are the first to show *Wolbachia* infections in *X. saxesenii* (85). Here we detected *Wolbachia* also from abandoned fungus gardens, which is surprising as these bacteria are obligate endosymbionts of insects (85). So either we were able to report a case of plant-mediated horizontal transmission, as it was already found for whiteflies (86) or the *Wolbachia* DNA originated from dead cells in the beetle’s faeces. The second hypothesis is supported by the lower abundances in the *removal* treatment for these lineages. The role of *Wolbachia* for *X. saxesenii* remains enigmatic as no obvious effects on beetle fitness or sex ratio could be detected in the few infected families. However, our bacterial communities were either dominated by *Pseudoxanthomonas* or *Wolbachia*, which could either be caused by an indirect effect through the host (e.g. host genome differences or an upregulation of immunity as response) or just sequencing dominance by *Wolbachia*.

### The fungal community and potential functions

The core community of fungi is small and made up of *C. globosum* (Chaetomiaceae), *R. sulphurea, R. canadensis* (Ophiostomaceae), *P. commune* (Trichocomaceae) and *A. biseptum* (Bionectriaceae). Among the secondary symbionts, *C. globosum* was the only species present in all nests. It is a saprophyte that is in obvious competition with the two *Raffaelea* ambrosia fungi. The same nature of competing interaction may be the case for *A. biseptum* as indicated by the replacement of the food fungi in some nests (Fig. S2). *C. globosum* is commonly isolated from wood and wood-boring insects, and it is also known to be toxic to insects (87). Its negative effect on beetles is further illustrated by the reduced success rate of 2^nd^ founding attempts when it was common in removal nests (Fig. S6B). However, beetles seem to have some strategies to control the spread of this fungus as demonstrated by the reduced abundance when mothers and immatures are present (Fig. 3D). *Penicillium* and *Acremonium* species may be regarded as weak competitors of the food fungi as they especially dominate in old nests (40,88). All of these secondary fungi are ubiquitous saprophytes within wood, do not infect beetles and are among the most common secondary symbionts of bark and ambrosia beetles (89).

### Strong heredity of beetle microbiome

Apart from the effect of farming there were strong signatures of family lineage on microbial symbiont communities between F0 and F1 generations. This means that the diversity and abundance of both cultivars and secondary symbionts was inherited over generations. For the cultivars that was expected, given that *X. saxesenii* females vertically transmit spores of their *Raffaelea* cultivars in elytral mycetangia and the gut (34). However, it is fascinating that this strong signature of family even prevails when females are removed from the nest and symbiont communities grow without the beetle’s presence (Fig. 3D, Fig. S2). The latter finding is a strong indication for the competitive abilities of the *Raffaelea* cultivars, which are apparently able to maintain their growth, possibly by producing antimicrobial compounds (e.g. ethanol; (90)). The in our data detected family-specific strain variation within *R. canadensis* is another clear sign for a strong heredity of symbionts. On the other hand, vertical transmission is also strong for the secondary symbionts. This may be surprising given that most of these taxa are likely competitors and pathogens of the mutualism. A closer look on these taxa reveals, however, that the majority are not only ubiquitous in bark and ambrosia beetles, but also many other saproxylic insects (91,92), and so even though we do not know much about the mechanisms, it seems that they are specialized for hitch-hiking and dispersal by insects.

### Usage of special primers for fungal metabarcoding helps to distinguish between food fungal species

Up to now there has been no satisfying method for metabarcoding of the fungal symbionts of bark and ambrosia beetles, as the universal, highly variable ITS primers used for species identification of fungi do not amplify the Ophiostomataceae, their primary group of symbionts (37,93). In line with Skelton et al. (51) and Ibarra-Juarez et al. (36), our work is another attempt to use self-designed non-ITS primers for amplicon sequencing of the fungal symbionts of an ambrosia beetle. Originally designed for this study, our LSU primer pair LIC15R and nu-LSU-355-3’ has already been used in another recently published article by Nuotclà et al. (38). Both of our studies successfully amplified the ambrosia fungi in the Ophiostomataceae. Despite of a mean amplicon length of 276.74 bp, we achieved a reliable discrimination of closely related species, which is specifically important for the two *Raffealea* ambrosia fungi of *X. saxesenii* (*R. sulphurea* and *R. canadensis*). Likewise, it was possible to identify the majority of other fungal symbionts, which included the following ascomycete orders: Ophiostomatales, Eurotiales, Sordariales, Hypocreales, Capnodiales, Onygenales, Helotiales, Xylariales and Dothideales. Yeasts of the order Saccharomycetales, however, that were successfully amplified by the SSU primers used by Ibarra-Juarez et al. (36) failed to be detected by our primers as we could demonstrate in the mock community control output. For future studies that aim to comprehensively cover bark and ambrosia beetle fungal symbionts, we propose a combined use of LSU ((51); this study) or SSU primers (36) with commonly used ITS primers (e.g. ITS3 & ITS4 or ITS1F & ITS2; (94,95)).

## Conclusion

In this study we tested the presence of active farming behaviour in ambrosia beetles. A significant effect of mothers and their larvae on the microbial composition of their fungus gardens could be proven and even stronger farming effects of adult daughters at a later developmental stage of nests may be expected. However, the exact mechanisms underlying the defence against weeds and the potential promotion of ambrosia fungi still remains unknown. Most studies investigating microbial symbionts of ambrosia beetles are surveys and we need more experimental studies on symbiont communities to understand the roles of specific symbionts in these ecosystems. Our new amplicon primers, which amplify Ophiostomatacea symbionts of bark and ambrosia beetles could distinguish the two ambrosia fungi in this study and therefore provide a useful tool for this purpose. In combination with more advanced techniques like quantitative real-time PCR this should help to understand symbiont shifts under different conditions. Also, we can move on to closely investigating some of the core taxa that we found (*Pseudoxanthomonas, Wolbachia, Penicillium*) in bioassays to find out more about their potential functions in the ambrosia beetle-fungus mutualism.

## Acknowledgements

We thank Mr. Schönmüller of the Würzburg forestry operation for the beetle collection permit.

## Funding Statement

This project was funded by the German Research Foundation (DFG) (Emmy Noether grant number BI 1956/1–1 to PB).

## Conflict of Interest

The authors declare that the research was conducted in the absence of any commercial or financial relationships that could be construed as a potential conflict of interest.

## Notes

https://github.com/janinad88/ambrosia-beetle-fungus-farming

### Competing Interest Statement

The authors have declared no competing interest.

