## Supplementary Material for "First experimental evidence for active farming in ambrosia beetles and strong heredity of garden microbiomes"

### Sequencing controls

To ensure the reliability of our symbiont community results, we included bacterial mock communities, fungal mock communities and negative controls. The "Microbial Community Standard" from ZymoBIOMICS was used as a bacterial mock community. Our fungal mock community consisted of five known fungal associates of *X. saxesenii*. The two primary symbionts, *R. sulphurea* and *R. canadensis*, the contaminant *Chaetomium globosum* as well as *Sporothrix stenocerans* and the yeast *Pichia sp.*.

To further enhance the quality of our data, we ran the contaminant removal method with the R package 'decontam' (1), taking into account the 'negative' control samples (*n* = 4; autoclaved rearing medium for beetle breeding and PCR water control) which had overall a very low read number of bacterial and fungal ASVs. This filtering process reduces the complexity of microbiome data while preserving their integrity in downstream analysis. A reduction of the classification methods' sensitivity and of technical variability, allows the generation of more reproducible and comparable results in microbiome data analysis (2).

### Bioinformatic processing and reference databases

The raw sequence reads were obtained from the Illumina MiSeq output directly (sample reads already demultiplexed by the MiSeq Reporter v. 2.5.1.3 with perfect index matches only). We merged the forward and reverse reads using the *-fastq_mergepairs* command in USEARCH v11.0-2.667 (3). A first quality filtering (sequence length >200 and maximum differences in the alignment = 30) was performed and overall quality was assessed with the command –*fastq_filter* specifying a high expected error threshold of 1 (*fastq_maxee*). Unique sequences in our FASTq file were identified with the command *–fastx_uniques* and sorted by decreasing size annotation with a minimum size of 4 (*-sortbysize*). Before clustering, we denoised the amplicon reads with *–unoise3* of the UNOISE algorithm. Instead of the traditional Operational Taxonomic Units (OTUs) our attempt produced ASVs (amplicon sequence variants, (4)) as a higher resolution option identifying biological sequences after denoising. With the –*usearch_global* command we set the identity threshold for the 16S taxa first to 99% identity.

ASVs were taxonomically classified in three steps with RDP classifier and the rdp_16s_v16_sp.fa reference database (5). In the first step, we used the reference database with manually added sequences of *Pseudomonas fluorescens* isolated from *X. saxesenii* in our laboratory. In a second step, all unclassified ASVs gathered in a "no hit"-file were classified with the rdp_16s_v16.fa database at 99% identity threshold and afterwards the remaining unclassified taxa with a sintax cut off from 0.8. Two reference databases were used for the classification of the LSU ribosomal RNA sequences (see also (6)): Firstly, we used a LSU rRNA sequences database of a fungal stock culture at the Chair of Forest Entomology and Protection (University of Freiburg, Germany). This database from known fungal symbionts of ambrosia beetles included eighteen reference sequences of twelve unique fungal species. Since this database comprises all the sequences of known fungal symbionts of *X. saxesenii*, we lowered the identity threshold to 97% identity. Secondly, ASVs not receiving a hit were classified with a custom reference database of fungi from NCBI, created with BCdatabaser v.1.1.1. (7), using the usual identity threshold of 99% identity and if still classifiable afterwards hierarchically with USEARCH Sintax using a cut off of 0.8. The newly created reference database from NCBI sequence data included 85,250 sequences fungal species (https://zenodo.org/record/5109344#.YPFDjEBCSUk). Finally, all tables, each for 16S and LSU taxonomic data, were combined into a common table. For a full script on parameter settings of all steps above, see the supplemented Shell-scripts in the GitHub repository.

**Overall symbiont communities**

In total we obtained 36,625,164 raw 16S reads and 24,009,500 raw LSU reads, which accounted for an average of 95,378 reads for 16S and 62,525 reads for LSU per sample in our two sequencing runs including several other projects. The total raw read size for this specific project was 1,175,496 raw 16S reads and 2,008,472 raw LSU reads, which accounted for an average of either 20,623 or 35,236 reads per sample. After data processing (merging, low quality <Q20, short reads <200 bp, ambiguous base-pairs), a mean of 18,056 reads per 16S sample and 16,995 per LSU sample remained. After read processing, and removing nonbacterial/fungal and rare sequences (<500 reads across sample set), the bacterial samples without community standard and negative controls contained on average 15,012 reads (range, 793 to 44,924) and 69 amplicon ASVs. The fungal samples contained on average 17,344 reads (range, 1087 to 39,932) and 202 amplicon ASVs.

Negative controls (autoclaved rearing medium for beetle breeding and PCR water control) showed very few bacterial (< 80 reads/sample) and fungal reads (< 220 reads/sample). In addition, the first controls of these samples with gel electrophoresis ahead to sequencing revealed no visible bands and rarefaction curves as well as richness estimates suggest a low number of single sequences due to possible cross-contamination (Supplementary Figures 1).

Positive mock controls for both bacteria (ZymoResearch, ZymoBIOMICS Microbial Community Standard D6300) and fungi data (self-designed community consisting of equal ratios of cultivated *C. globosum, R. sulphurea, R. canadensis, Sporothrix stenocerans* and *Pichia sp.*) confirmed successful sequencing of the contained taxa. However, the yeasts (*Pichia sp., Saccharomyces cerevisiae* and *Cryptococcus neoformans*) from the ZymoBIOMICS and fungal mock control could not be detected with the LSU primers we used. Bacterial species from the ZYMO mock community rarely appeared in other samples than the mock communities representing neglectable cross-contamination (<15 reads/sample).

The contaminant removal method identified 178 of the bacterial 16S ribosomal RNA and two of the fungal LSU ribosomal RNA ASVs as external contaminants, which were excluded from the data. Species accumulation curves of the final data sets showed that most samples were sequenced to saturation after approximately 20,000 high quality reads for 16S (Fig. S1a). The rarefaction curve for LSU sequence data reached the plateau after approximately 5,000-6,000 reads (Fig. S1b).

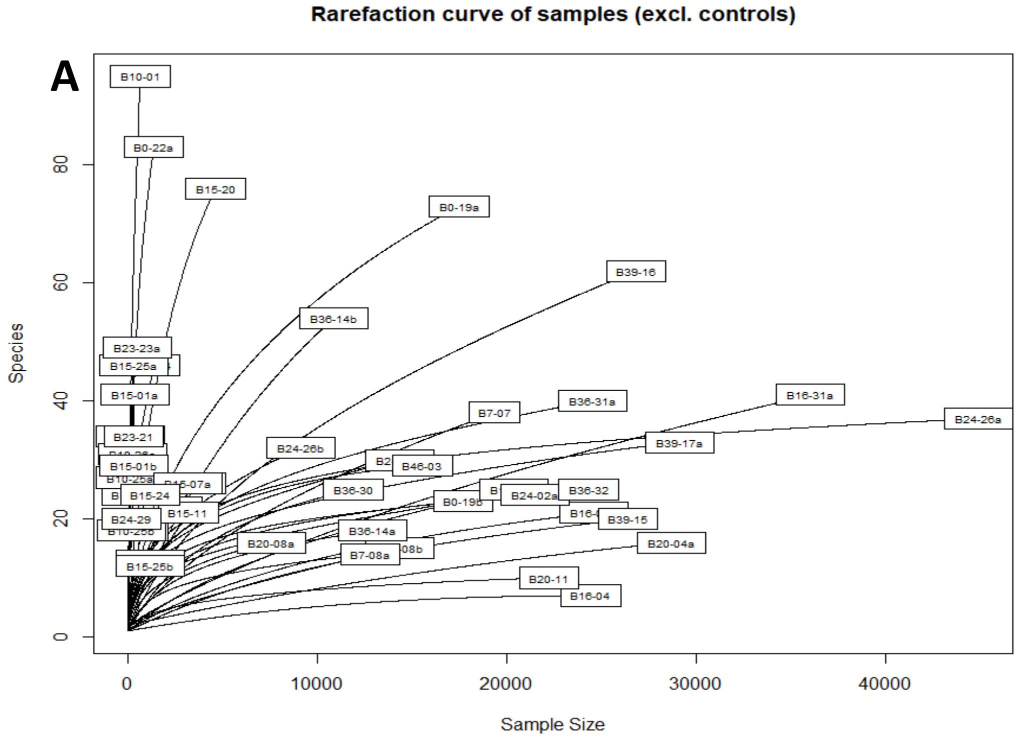


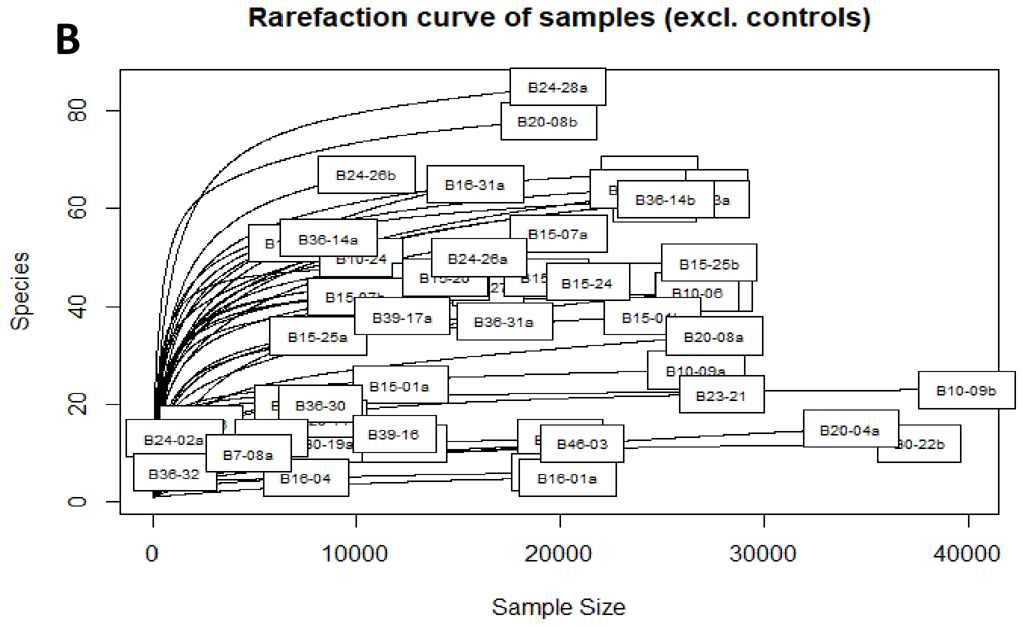


**Fig. S1: rarefaction curves for 16S (A) and 28S (B) amplicon sequence variants in the final datasets.**

**Tab. S1: Relative abundance of the most abundant bacterial taxa in the three treatments (mean ± SD).**

| Genus | Treatment | Mean RA (%) |
|---|---|---|
| *Acinetobacter* | control | 0.57 ± 1.43 |
| | removal | 0.16 ± 0.57 |
| | 2nd attempt | 0.02 ± 0.03 |
| *Erwinia* | control | 0.29 ± 0.78 |
| | removal | 15.41 ± 32.43 |
| | 2nd attempt | 7.57 ± 20.75 |
| *Ochrobactrum* | control | 4.51 ± 16.90 |
| | removal | 1.88 ± 6.90 |
| | 2nd attempt | 0.25 ± 0.68 |
| *Pseudomonas* | control | 0.97 ± 3.35 |
| | removal | 0.28 ± 1.00 |
| | 2nd attempt | 0.08 ± 0.10 |
| *Pseudoxanthomonas* | control | 50.94 ± 47.03 |
| | removal | 67.85 ± 41.61 |
| | 2nd attempt | 54.78 ± 48.03 |
| *Streptomyces* | control | 0.80 ± 2.77 |
| | removal | 0.05 ± 0.14 |
| | 2nd attempt | 0.02 ± 0.05 |
| *Wolbachia* | control | 36.88 ± 44.08 |
| | removal | 11.55 ± 29.90 |
| | 2nd attempt | 36.15 ± 49.15 |

**Tab. S1: Relative abundance of the most abundant fungal species (>0.5%) in the three treatments (mean ± SD).**

| Species | Treatment | Mean RA (%) |
|---|---|---|
| *Chaetomium globosum* | control | 18.98 ± 28.11 |
| | removal | 28.75 ± 39.06 |
| | 2nd attempt | 25.37 ± 34.99 |
| *Raffaelea sulphurea* | control | 24.74 ± 29.65 |
| | removal | 15.37 ± 26.34 |
| | 2nd attempt | 12.66 ± 20.63 |
| *Raffaelea canadensis* | control | 5.52 ± 9.29 |
| | removal | 1.88 ± 4.37 |
| | 2nd attempt | 5.88 ± 10.64 |
| *Penicillium commune* | control | 0.001± 0.004 |
| | removal | 0.71 ± 3.17 |
| | 2nd attempt | 3.17 ± 10.51 |
| *Acremonium biseptum* | control | 0.20 ± 0.90 |
| | removal | 4.82 ± 21.9 |
| | 2nd attempt | 2.37 ± 7.85 |

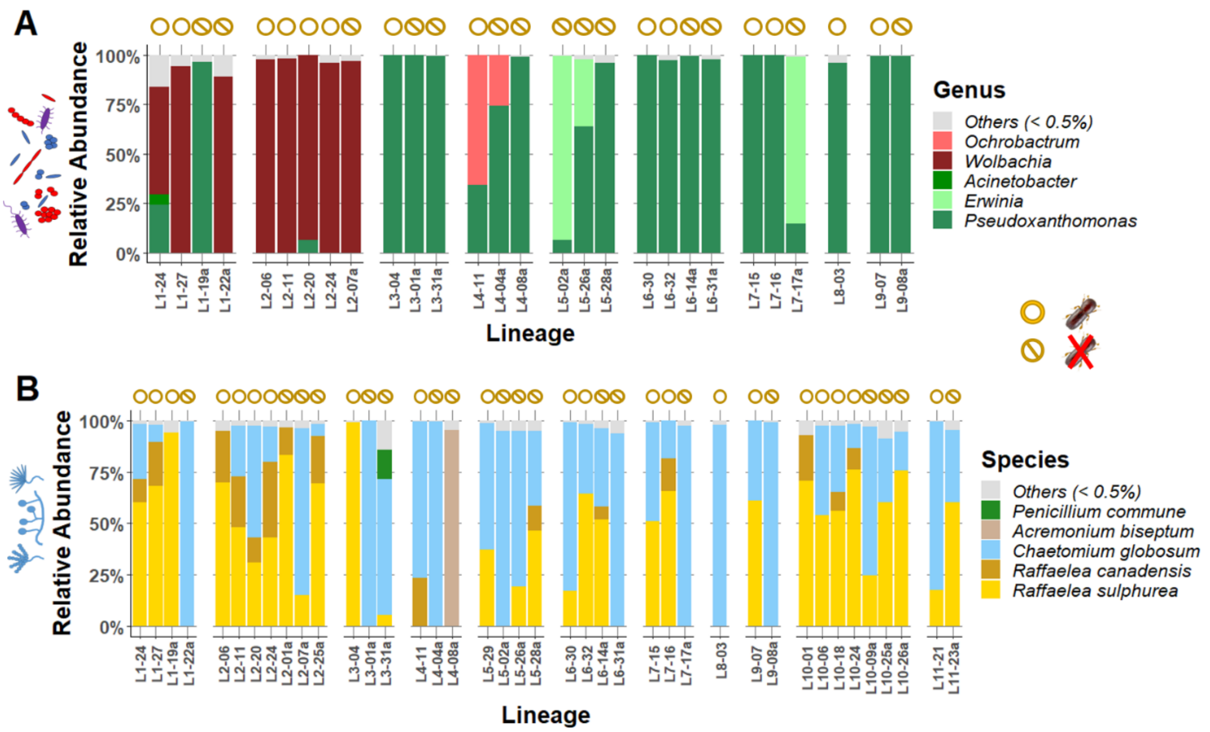


**Fig. S2: Effect of family lineage on microbial communities.** The core bacterial (A) and fungal community (B) of *X. saxesenii* is small. In addition to the treatment effect there is also a significant effect of family lineage on both bacterial and fungal communities (for details see Tab. 1). Only bacterial and fungal taxa with a relative abundance of >0,5% are displayed (everything else is combined in "others"). *Control* nests are marked by closed circles, *removal* nests by crossed out circles.

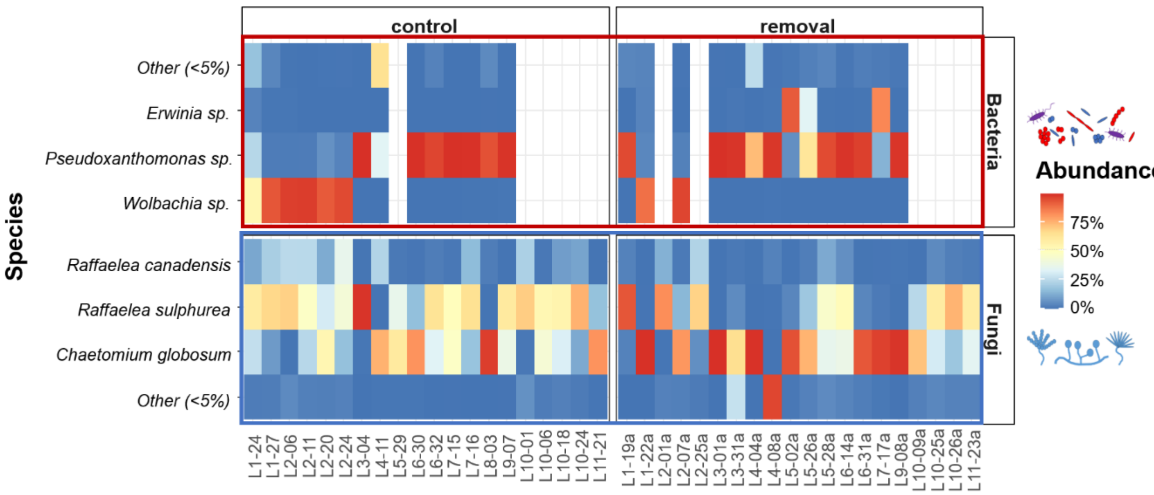


**Fig. S3: Heatmap of relative abundance of core bacterial and fungal phylotypes in all samples displayed for *control* and *removal* group.** Taxa under the detection threshold of 5% relative abundance are combined into "Other".

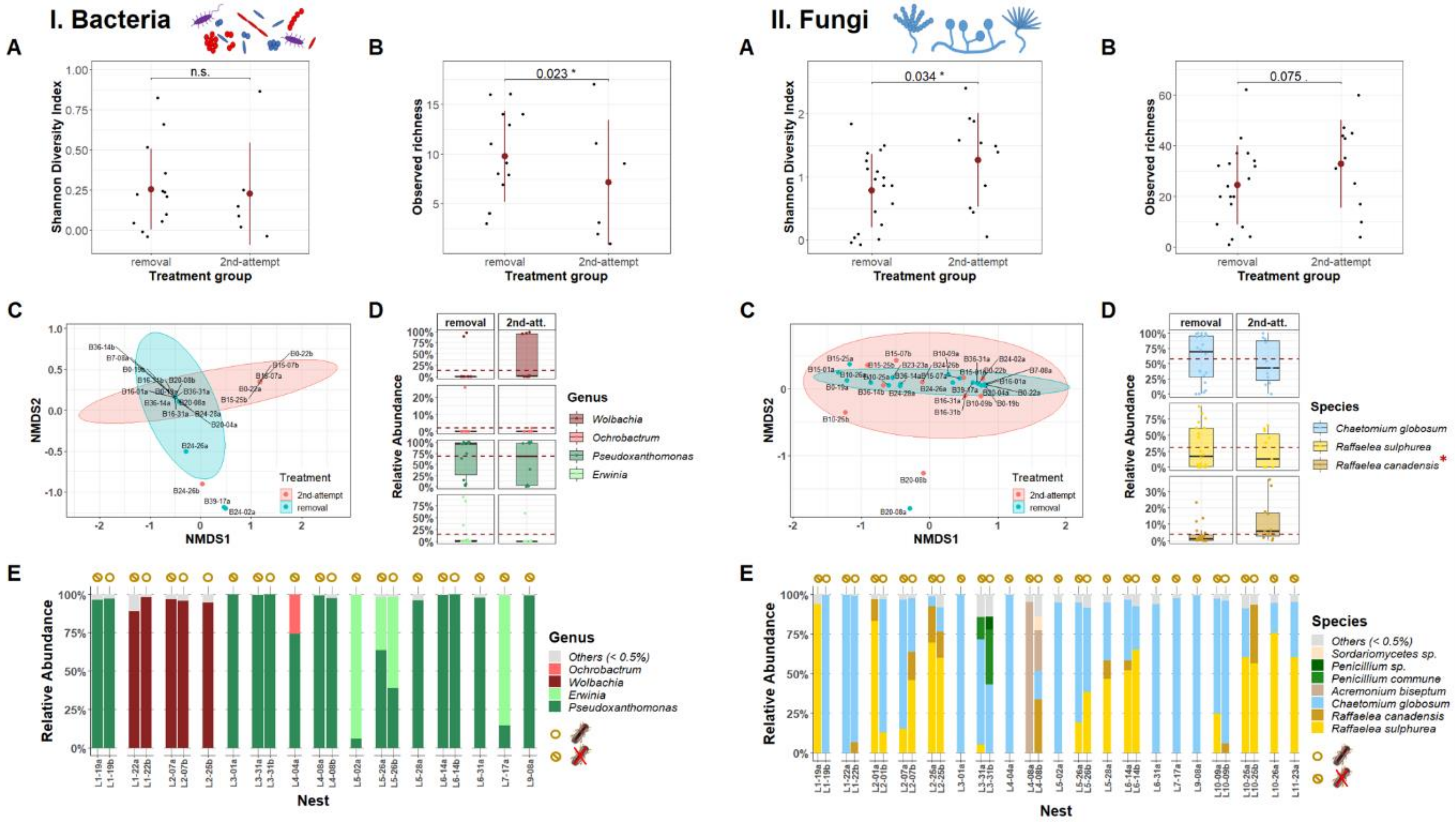


**Fig. S4: Effect of beetle removal on bacterial and fungal diversity, composition and abundance.** (A) Bacterial Shannon's diversity indices did not differ between *removal* ($n$ = 20 nests) and *2nd attempt group* ($n$ = 11 nests; LM: $F$ = 0.406, $p$ = 0.538), opposite to the fungal SDI, which was higher in the *2nd attempt group* (GLM: $\chi^2$ = 1.28, $p$ = 0.258; plots show mean and standard deviation). (B) Bacterial and fungal observed richnesswas higher when beetles were removed (Bacteria: GLM: $\chi^2$ = 4.51, $p$ = 0.034; Fungi: GLM: $\chi^2$ = 3.17, $p$ = 0.075; plot shows mean and standard deviation). (C) Visualization of compositional differences between bacterial and fungal communities of nests in *removal* and *2nd attempt group* (NMDS on Bray-Curtis dissimilarity: $stress_{Bacteria}$ = <0.001, $stress_{Fungi}$ = 0.032; ellipses represent 95% confidence intervals; labels represent family lineage of samples). PERMANOVA of bacterial and fungal communities showed no effect by beetle presence (Bacteria: $R^2$ = 0.042, $p$ = 0.169; Fungi: $R^2$ = 0.016, $p$ = 0.641). D). Comparisons of relative abundances of core bacterial taxa did not reveal any significant differences between treatments (boxplots represent median with its interquartile range and whiskers; red dashed line shows the mean relative abundance of the `*removal group*`). Comparisons of relative abundances of core fungal taxa revealed a reduction of the food fungus, *R. canadensis* (LM: $F$ = 4.30, $p$ = 0.051; * - $p$ < 0.05), in nests without beetles. E) The core bacterial and fungal community of *X. saxesenii* is small. There is a significant effect of family lineage on the bacterial community (for details see Tab. 1). Only bacterial and fungal taxa with a relative abundance >0,5% are displayed (everything else is combined in "others"). *Control* nests are marked by closed circles, *removal* nests by crossed out circles.

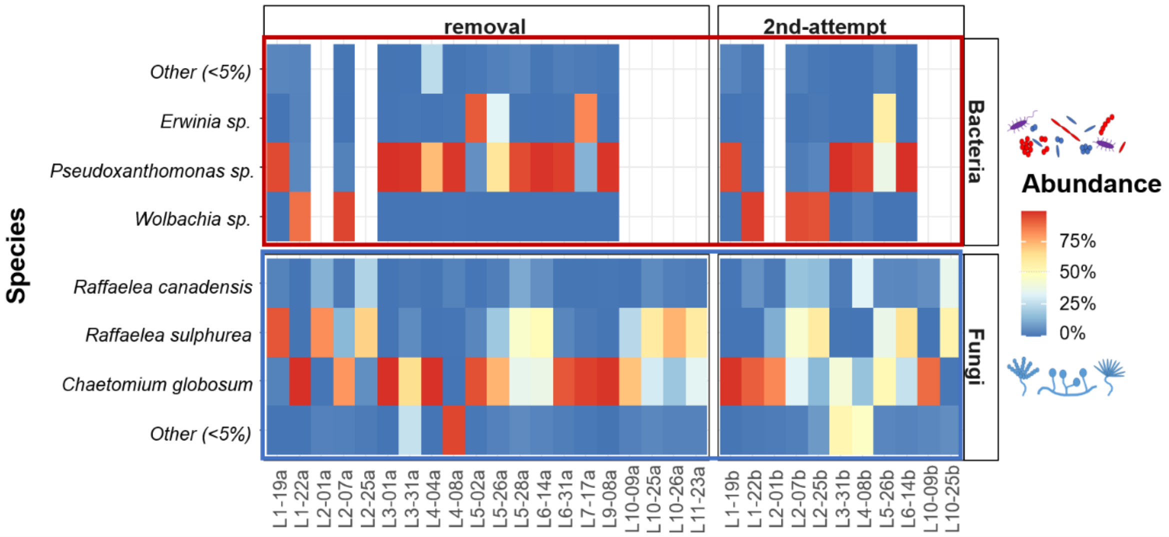


**Fig. S5: Heatmap of relative abundances of core bacterial and fungal phylotypes in all samples displayed for *removal* and 2$^{nd}$ *attempt* group.** Taxa under the detection threshold of 5% relative abundance are combined into "Other".

**Tab. S3: Statistical output of linear models on relative abundances of core bacterial and fungal taxa.** Both comparisons of *control* vs. *removal* and *removal* vs. 2$^{nd}$ *attempt* are shown. Statistically significant results are depicted in bold.

| ***control* vs. *removal*** | | | | | |
|---|---|---|---|---|---|
| **Bacteria** | | | **Fungi** | | |
| | *p*-value | | | *p*-value | |
| *Wolbachia* | Treatment | 0.138 | *Chaetomium globosum* | Treatment | 0.077 |
| | Lineage | **<0.001** | | Lineage | 0.618 |
| *Pseudoxanthomonas* | Treatment | 0.679 | *Raffaelea canadensis* | Treatment | **0.031** |
| | Lineage | **0.001** | | Lineage | **0.038** |
| | | | *Raffaelea sulphurea* | Treatment | **0.034** |
| | | | | Lineage | **0.095** |
| ***removal* vs. 2$^{nd}$ *attempt*** | | | | | |
| **Bacteria** | | | **Fungi** | | |
| | *p*-value | | | *p*-value | |
| *Wolbachia* | Treatment | 0.534 | *Chaetomium globosum* | Treatment | 0.765 |
| | Lineage | **0.045** | | Lineage | 0.562 |
| *Pseudoxanthomonas* | Treatment | 0.693 | *Raffaelea canadensis* | Treatment | **0.051** |
| | Lineage | **0.005** | | Lineage | 0.623 |
| | | | *Raffaelea sulphurea* | Treatment | 0.241 |
| | | | | Lineage | **0.051** |

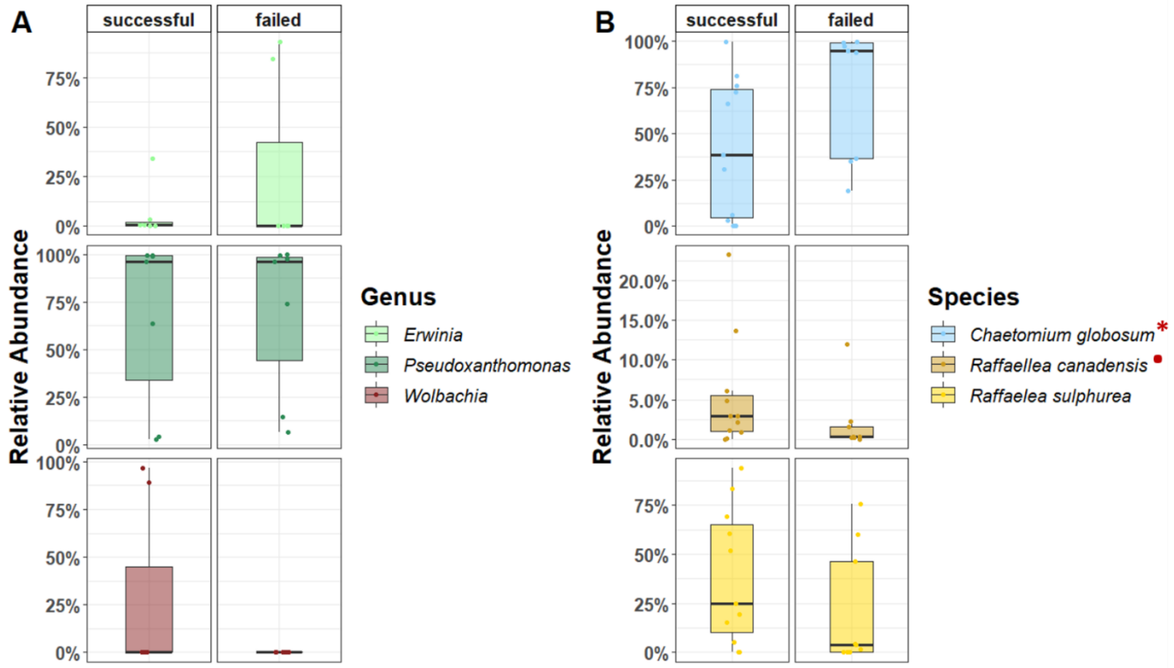


**Fig. S6: Effects of the relative abundance of core taxa in *removal* nests on the success of the same foundresses in their 2nd nest-founding attempt.** There was no significant effect of core bacterial taxa on founding success, but relative abundance of *C. globosum* correlated with a lower founding success (* - $p < 0.5$), while *R. canadensis* tended to correlate with a higher founding success (• - $p = 0.1$) (boxplots represent median with its interquartile range and whiskers).

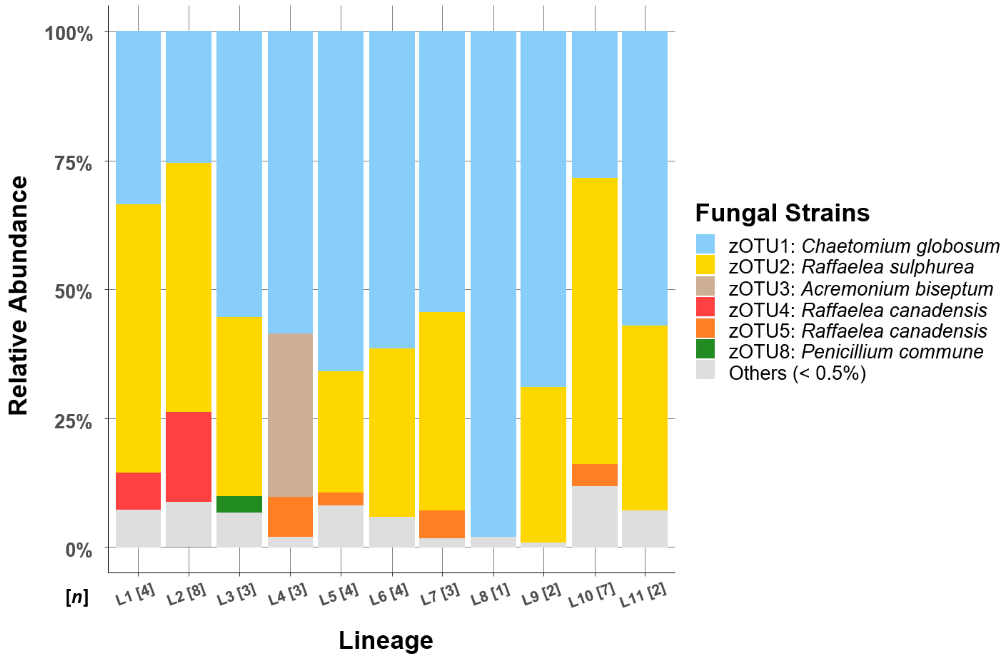


**Fig. S7: Heritability of fungal communities in control nests.** Visualization of highly abundant strains in ambrosia beetle nests, revealed the presence of two *R. canadensis* strains each only individually dominant per family lineage.
